# Repertoire of morphable proteins in an organism

**DOI:** 10.1101/719260

**Authors:** Keisuke Izumi, Eitaro Saho, Ayuka Kutomi, Fumiaki Tomoike, Tetsuji Okada

**Affiliations:** Department of Life Science, Gakushuin University, 1-5-1 Mejiro, Toshima-ku, Tokyo 171-8588, Japan.

**Author notes:** Corresponding author: Tetsuji Okada, Department of Life Science, Gakushuin University, 1-5-1 Mejiro, Toshima-ku, Tokyo 171-8588, Japan.

## Abstract

All living organisms have evolved to contain a set of proteins with variable physical and chemical properties. Efforts in the field of structural biology have contributed to uncovering the shape and the variability of each component. A set of experimental coordinates for a given protein can be used to define the “morphness/unmorphness”. Here we show the results of global analysis of more than a thousand *E. coli* proteins, demonstrating that it would be a comprehensive method of understanding the evolved repertoire in an organism. By collecting “UnMorphness Factor” (UMF) determined for each of the proteins, the lowest and the highest boundaries of the experimentally observable structural variation are understood. The distribution of UMFs obtained for an organism is expected to represent how rigid and flexible components are balanced. The present analysis extends to evaluate the growing data from single particle cryo-electron microscopy, providing valuable information on effective interpretation to structural changes of proteins and the supramolecular complexes. The data and the method presented here also conform to FAIR data principles, having potential significance to advance the field of structural and molecular cell biology.

## Introduction

Self-replicating (living) species are defined by the presence of a genome that is used to produce a set of proteins and RNAs. The diversity of living entities, from unicellular bacteria to mammals, primarily corresponds to the varying repertoire of proteins. A proteome set limits the morphology and functionality of an organism. Therefore, understanding the atomic details of the folded state and the variability of the fold for each protein in an organism is a long-term goal in structural biology.

We have previously shown that comparison of different proteins and protein families with respect to their structural variability can be comprehensively performed by distance scoring analysis (DSA), in which all intramolecular C_α_-C_α_ pair distance variations (score = average / stdev) are considered^1^ (Fig. 1). DSA is advantageous in terms of reproducibility because no procedures/assumptions associated with superimposition are involved. Also, the amount of numeral usage is far more than the pair-wise based analysis. For example, if we calculate an average root-mean-square deviation (RMSD) of C_α_ positions for 10 structures (chains) of 100 amino acids, the number of used deviation values is 100 positions × 45 structure pairs = 4,500; whereas, DSA uses (100 × 99 /2) C_α_-C_α_ pair distances × 10 structures = 49,500 scores. Furthermore, inclusion of new data is simpler in DSA as no pairing is required.

**Fig. 1.**
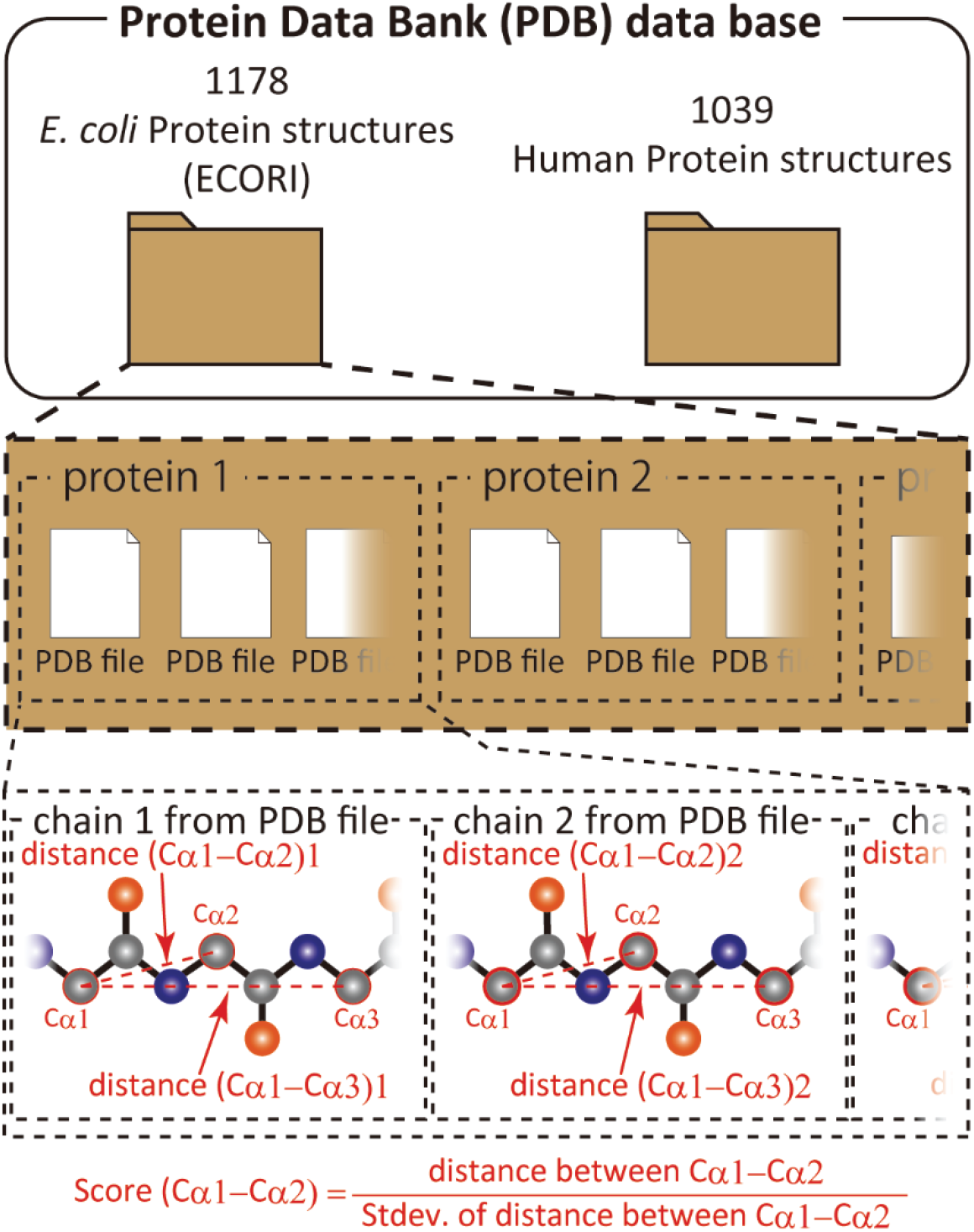
Scheme of score calculation.

*E.coli* proteins are the most abundant prokaryotic content in the Protein Data Bank (PDB)^2^. We propose that species-based construction of a “morphability” map is most achievable, covering a substantial portion of proteins that construct the *E. coli* proteome. In this work, we demonstrate that a substantial fraction (~23%) of the *E.coli* proteome can be described as exhibiting “morphness/unmorphness” distribution with well-defined lower/upper limits, and a major cluster of data points. The result is compared to a similar amount (~1000) of randomly selected human proteins, providing unprecedented insight into the morphness repertoire for a set of natural polypeptides. These analyses of X-ray structures are also used to evaluate the recently accumulating cryo-EM structures.

## Results and discussion

### UMF, a comprehensive measure of protein structure variability

We previously proposed that “UnMorphness Factor” (UMF), which is defined as a converged average score for a protein, is an alternative to conventional RMSD-based comparison^3^. UMF is calculated with at least three chains of continuously modeled C_α_s. In most cases, UMF converges as the number of used chains increases. In the following study, UMFs calculated from X-ray crystallography are denoted as xUMF and those from single particle cryo-electron microscopy (cryo-EM) are denoted as eUMF.

Since many proteins are deposited to the PDB from a variety of *E. coli* strains, we chose a reference proteome set called ECOLI (strain K-12) in UniProt^4^. According to UniProt reference proteome sets, the proteome size ranges from 4232 (strain 333) to 6593 proteins (strain NCTC13148). Thus, the number of proteins in the ECOLI reference set (4391) is likely to be close to the minimum number necessary for constituting this prokaryotic organism.

As of July in 2019, the PDB contains over 8500 X-ray entries that include *E.coli* components (mostly proteins). We calculated xUMFs for all *E. coli* strains, totaling 1178 proteins, including redundancy in the case of some proteins. An example of this protein redundancy is β-lactamase, for which many UniProt IDs are found in the various strains of *E. coli*. Thus, 1178 xUMF values were mapped to the reference proteome set ECOLI, resulting in 980 proteins and 22.3% (980 / 4391) coverage of the whole reference proteome. On the other hand, when we re-calculated the coverage separately for soluble and membrane proteins assigned by SCAMPI^5^, a significant difference was found; 26.9% (887 / 3300) for soluble proteins and 8.5% (93 / 1091) for membrane proteins. Thus, our present analysis provides xUMFs over a quarter of soluble proteins in *E. coli* K-12. The average ratio of the analyzed to the full length (including signal/propeptide sequences) for 980 proteins is 83.6% (only 6.7% of 980 proteins is limited to <50% of full length).

For comparison, we have updated the previous human protein analysis and collected 1039 xUMFs. The SCAMPI analysis for the reviewed human reference set of 20,416 proteins assigned 6293 as membrane proteins. Since 221 out of 1039 xUMFs are of the possible 6293 membrane proteins, the coverage is 3.5%; whereas, 818 out of 14123 (=20416-6293) corresponds to 5.8% coverage for human soluble proteins.

In this study, proteins are selected for DSA when at least 3 chains (for *E. coli*) or 3 entries (for human) are available, with an exception for the case that the continuously modeled part is too short against the full length. Overall, the average usage per protein is 7.9 entries (10.1 chains) for the ECOLI reference set and 13.9 entries (15.4 chains) for the human proteins. The higher number in the human set is due to the data abundance in the PDB and to several heavily deposited proteins (such as carbonic anhydrase 2, for which we have used over 600 entries to calculate the xUMF). When we recalculate the numbers for the human set after removing the 12 proteins for which more than 100 entries/chains are used, the average usage drops to 10.9 entries (12.5 chains). For the ECOLI set, only maltose-binding periplasmic protein exceeds 100 entry/chain usage. These considerations indicate that the overall amount of data usage is not very different between the ECOLI set and the human protein set. On the other hand, a substantial portion of the ECOLI set xUMFs originates from the analysis of using only one or two entries per protein, taking multiple chains per entry. This limitation is presumably reflected in the xUMF distribution plot as shown below in Fig. 3.

### Summary plot for *E. coli* and human proteins

When all 980 xUMFs of the ECOLI set proteins are plotted against the analyzed chain length in log-log scale, a fairly dispersed image can be obtained (Fig. 2a). This is a log version of the previously described “summary plot”^1^. A corresponding figure for 1039 human proteins is also shown in Fig. 2c. These two images are similar despite all proteins being chosen randomly and only by giving selection priority based on the availability of the experimental coordinates by X-ray crystallography. An obvious difference between the ECOLI set and the human data is the amount of high xUMF proteins in the former and this presumably occurs because xUMFs of a substantial part of the ECOLI set proteins are calculated using less than three PDB entries; this means that more than one chain per entry had to be used. In fact, of the 38 proteins exhibiting xUMFs larger than 500, 34 proteins are analyzed using only one or two entries.

**Fig. 2.**
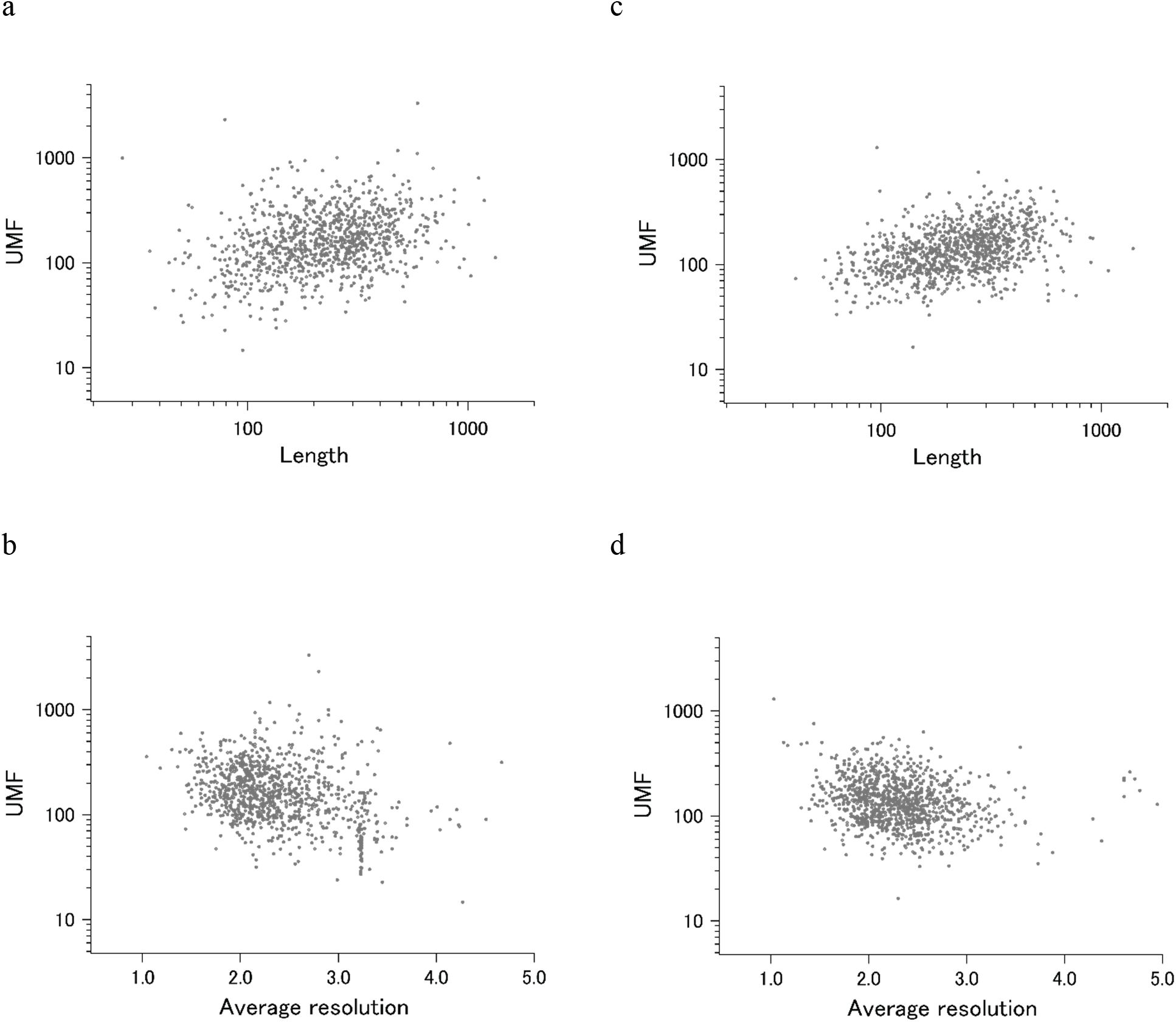
The summary plot of xUMFs. (a) log-log plot of ECOLI set against analyzed chain length; (b) log-log plot of human proteins against analyzed chain length; (c) semi-log plot of ECOLI set against average resolution (Å); (d) semi-log plot of human proteins against average resolution.

When the horizontal axis is changed to the average resolution, a cluster of proteins becomes clearer. In the ECOLI plot (Fig. 2b), ribosomal proteins, which includes thirty-one 50S and twenty 30S subunit proteins analyzed using mostly the same set of PDB entries, are shown as a thin cluster at the average resolution from 3.2 to 3.3 Å and the xUMF range from 25 to 70. On the other hand, COP9 signalosome complex subunits dominate the data of average resolution lower than 4.5 Å in the human summary plot (Fig. 2d). Overall, xUMFs appear to exhibit only weak dependence either on the polypeptide length or on the X-ray data resolution (correlation coefficients of 0.19, −0.12, 0.30, −0.21 for panel a, b, c, d, respectively). We have previously reported correlation coefficients corresponding to Fig. 2d of the present study but for ~100 proteins × 3 sets separated according to the amount of entry usage. The previous value of −0.33 (an average of 3 sets) decreased substantially to −0.21 for the 1039 proteins analyzed here. As a reference, similar analysis of over 5500 proteins from various species (including *E.coli* and human) resulted in correlation coefficients of 0.28 and −0.22 vs chain length and average resolution, respectively (unpublished results). These studies are consistent with a conclusion that xUMFs are more dependent on the chain length than on the data resolution.

### Distribution plot

The summary plots of ECOLI and human proteins confirmed that the lower limit of xUMFs is ~15, as previously reported for human calmodulin. As long as we analyze X-ray structures, xUMFs in the range from 0 to 10 (over 10% average deviations for all intramolecular C_α_-C_α_ pair distances) are not likely to emerge. In fact, based on the over 5500 proteins from various species analyzed so far, the lower limit of ~15 is still valid (unpublished result). Many proteins that are reluctant to crystallize would contain very flexible parts. If such flexible parts dominate the entire length of a polypeptide in a given protein, as can be seen in many NMR depositions (ensembles), the UMF should approach to 0. Thus, many intrinsically disordered proteins^6,7^ can be understood to exhibit 0~10 UMFs that cannot be evaluated by X-ray crystallography.

To demonstrate the probable upper limit of xUMFs, distribution of xUMF values shown in the summary plot are counted from 0 to 900 with the slot of 30 (Fig. 3). Whereas some proteins (8 out of 980 *E. coli* proteins, and 1 out of 1039 human proteins) exhibited the xUMFs larger than 900 and not shown in the figure, the plot demonstrates that most proteins would rarely exceed the xUMF over 500. This result means that the averaged uncertainty (standard deviation) of all C_α_-C_α_ distances in a folded polypeptide chain determined by X-ray crystallography inevitably exceeds 0.2% (0.1 Å per 50 Å). A clear difference between the ECOLI set and human proteins in the distribution plot is the smoothness of the pattern. In addition, the peak fraction is clearer in the human plot (Fig. 3b), where over 20% of the 1039 proteins are in the xUMF range of 90 to 120, corresponding to 1.1% to 0.83% average uncertainty of all C_α_-C_α_ distances.

**Fig. 3.**
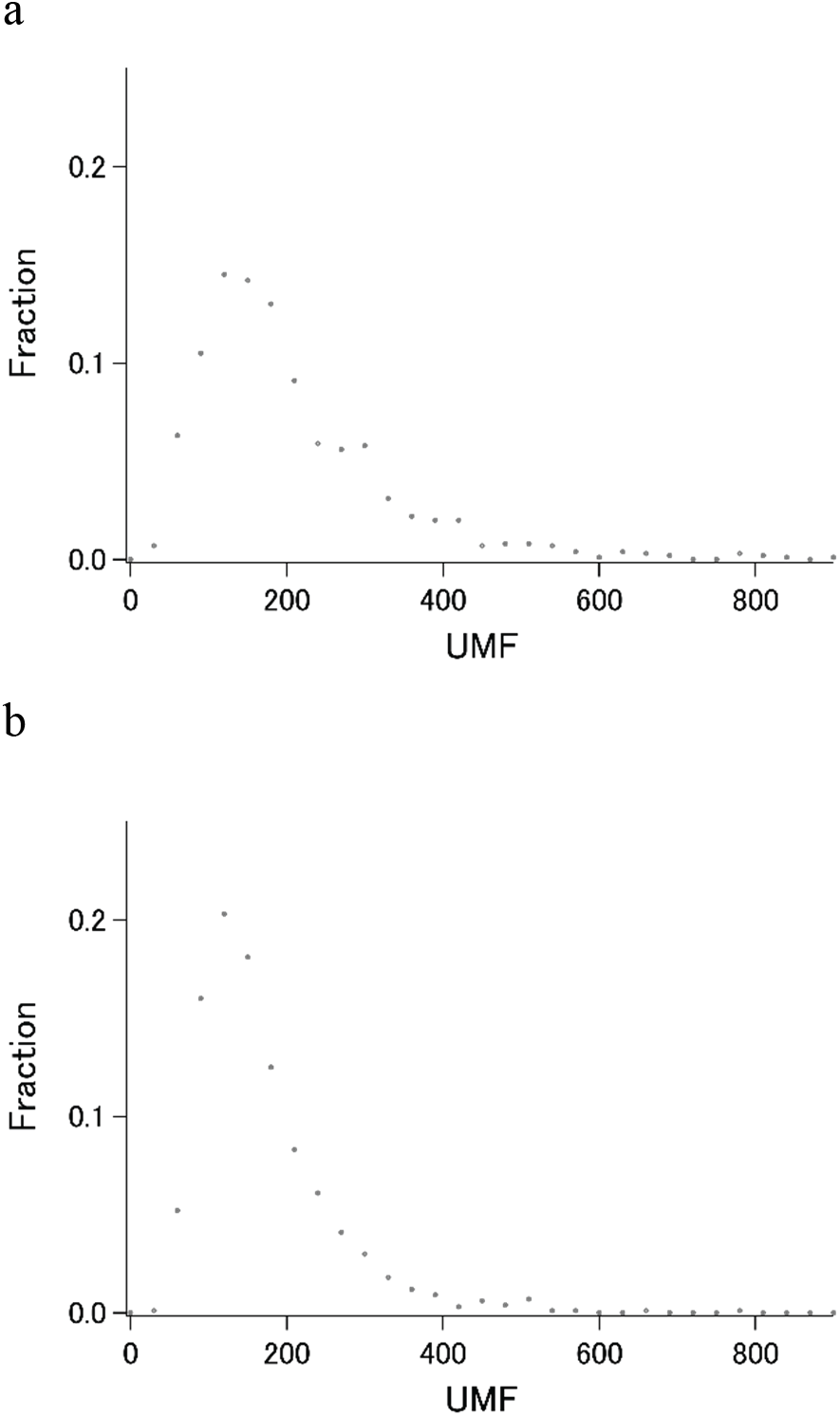
The distribution plot of xUMFs. (a) ECOLI set; (b) human proteins.

### Cryo-EM structure analysis

Cryo-EM data deposition is predominant in the PDB^8–10^. While the resolution of data by this method rarely reaches the precision that most X-ray structures can resolve, we have examined cryo-EM structures of ECOLI reference proteome set and calculated eUMFs for 56 proteins of which 31 have respective xUMFs. As a result, 1005 UMFs (980 xUMFs plus 25 eUMFs) out of 4391 ECORI set are obtained, corresponding to 22.9% coverage. Membrane proteins make up 11 of the 23 eUMFs, reflecting that cryo-EM is frequently applied to targets that are reluctant to crystallize.

When we add the eUMF data to the ECOLI reference set summary plot of xUMFs shown in Fig. 2, a clear distinction becomes obvious (Fig. 4). Importantly, eUMF can be seen more frequently in the long polypeptides and in the low value ranges, indicating that cryo-EM is effective in detecting conformational variability of large molecular weight proteins.

**Fig. 4.**
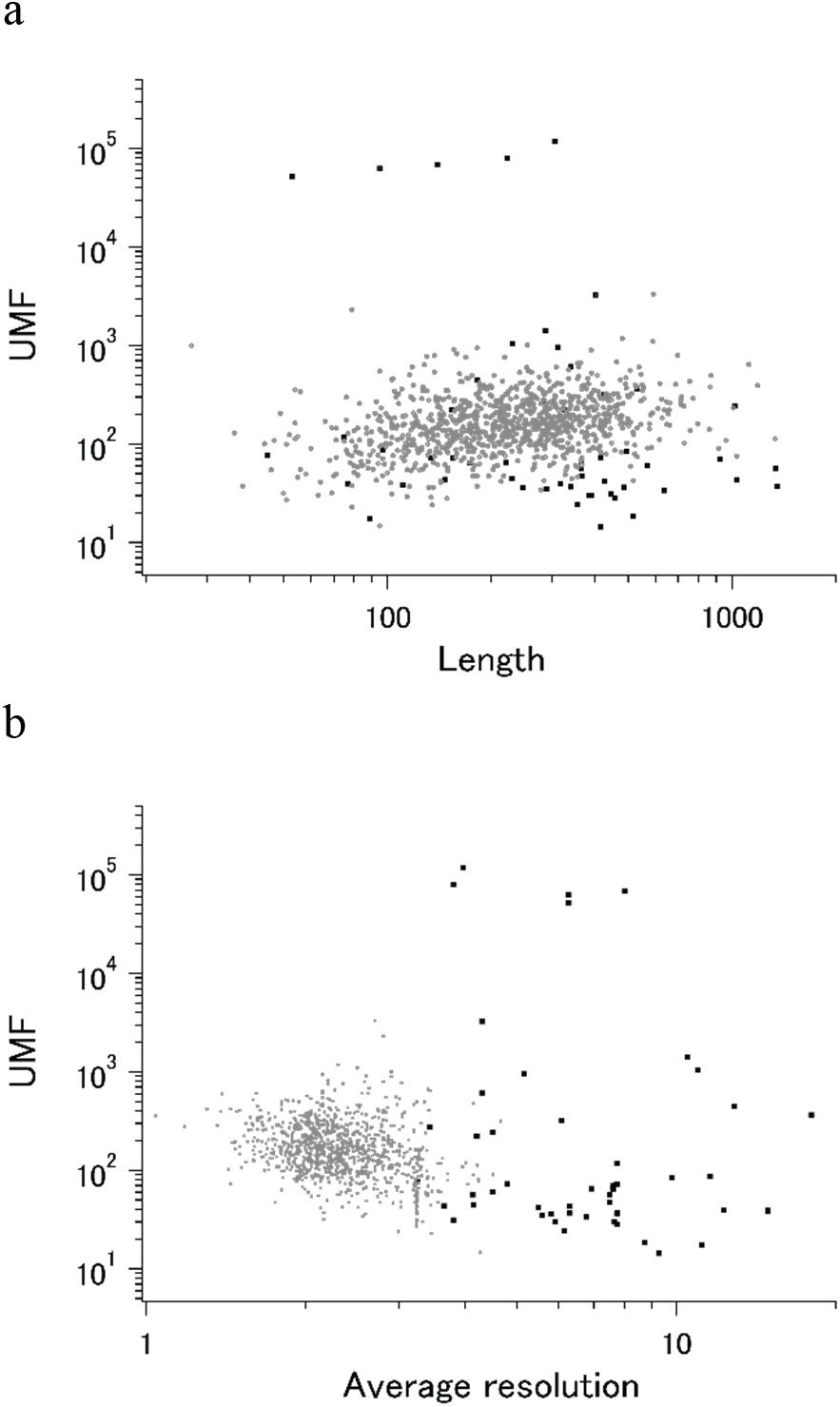
The summary plot of xUMFs and eUMFs. (a) log-log plot of ECOLI set against analyzed chain length; (b) log-log plot of ECOLI set against average resolution. eUMFs are shown as black squares.

DSA analysis of cryo-EM structures also provided some examples that are distinct from that obtained from a larger set of X-ray entries. The semi-log main plot, which is another form of the previously reported main plot^1^ and is a 2D representation of highly variable parts in a protein, is especially favorable in demonstrating how cryo-EM and X-ray structures can explore the morphness of proteins. In Fig. 5a, a semi-log main plot is shown for the cryo-EM entry (PDB ID: 5Y4O) of low conductance mechanosensitive channel YnaI^11^. This entry contains seven chains of identical length (223 residues, 65% of the full length). The unusually high eUMF of 79821.6 on this plot is explained by a large part of the polypeptide being virtually identical among the seven chains and clearly separated from the lower score thin cluster representing the C_α_ pair distances in the transmembrane (TM) part. Obviously, the eUMF in this case is far from the expected value for this protein. Six other proteins are also outstanding in the unusual value of eUMFs (Fig. 4).

**Fig. 5.**
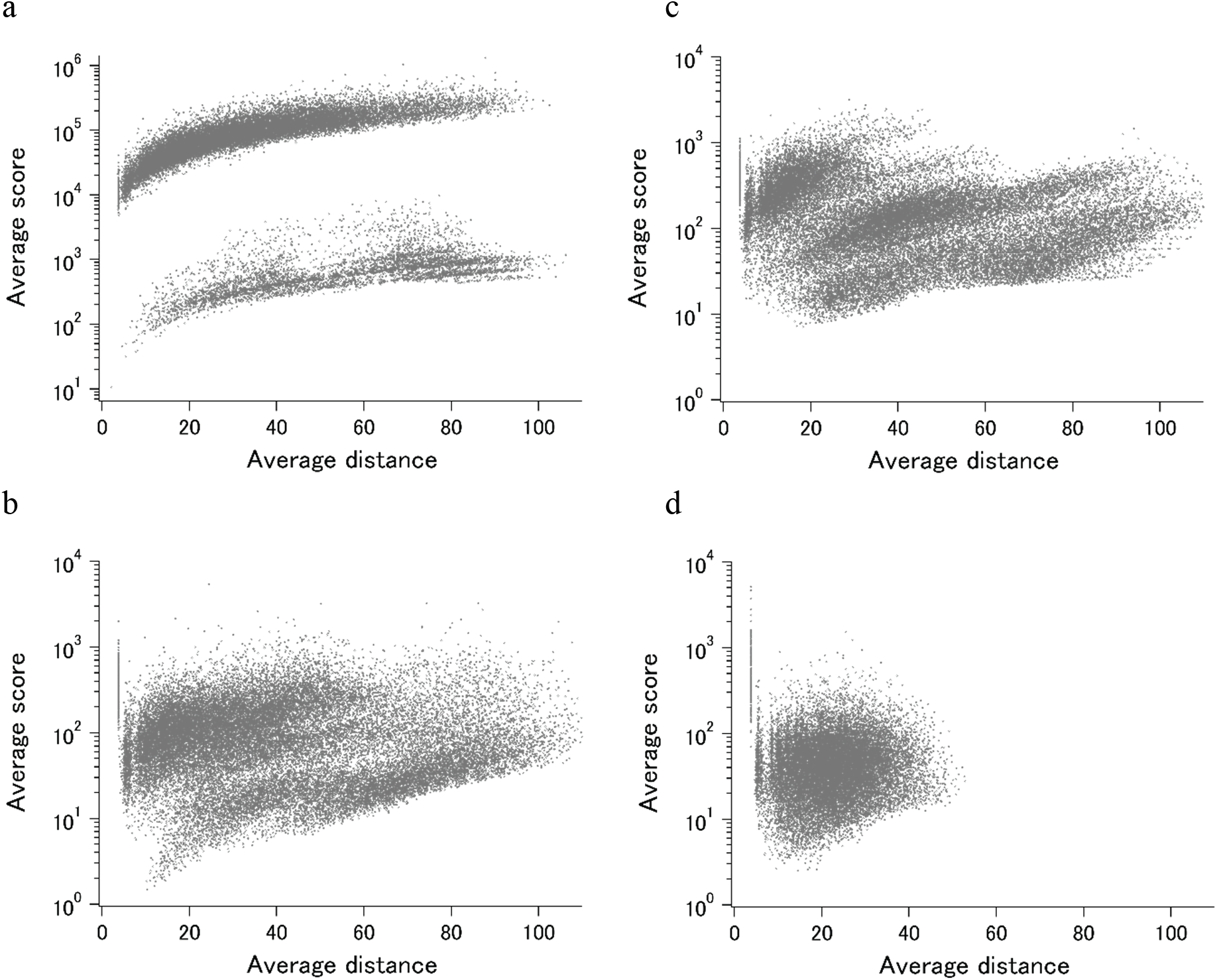
The semi-log main plot obtained from DSA. (a) YnaI of ECOLI (cryo-EM); (b) MscS of ECOLI by single entry analysis (X-ray); (c) MscS of ECOLI by multi entry analysis (X-ray); and (d) CNR1 of human (X-ray). The unit of average distance is Å.

YnaI is functionally related to the MscS mechanosensitive channel^12^. Although the sequence similarity between them is limited, they both self-assemble to form a heptamer, in which each protomer assumes an elongated conformation. Single entry analysis of MscS as has been done for YnaI, but using X-ray entry (PDB ID: 2OAU)^13^, resulted in xUMF of 190.6, which is considerably lower than eUMF of YnaI and fairly in the range shown in the distribution plot (Fig. 3a). The corresponding semi-log main plot demonstrated multidomain morphness (Fig. 5c), but not separated from each other in the average score range. Multi entry analysis (1 chain usage per entry and 6 entry usage) lowered the xUMF to 119.0, resulting in a diffused pattern on the semi-log plot (Fig. 5b). For comparison, the pattern of semi-log plot obtained for 200 residue TM bundle of a G protein-coupled Cannabinoid receptor 1 (CNR1) is shown in Fig. 5d. As the analyzed part is only limited to compactly arranged seven TM helices, the maximum dimension is about a half of YnaI and MscS. This receptor, whose conformational change occurs mostly in one of the seven helices^14^, is among many examples where any large domain morphness does not occur but the polypeptide contains very variable parts, resulting in a low xUMF (58.9 for CNR1).

For other two proteins, ribosome hibernation promoting factor (HPF) and ribosome modulation factor (RMF)^15^, out of the seven proteins with unusually high eUMFs (Fig. 4), there were respective X-ray entries (but not enough chains for xUMF calculation). By merging just one X-ray entry with the corresponding cryo-EM entries for each of the two proteins, the resulting UMF (xeUMF) dramatically decreased to 90.4 for HPF and 66.0 for RMF, falling into range on distribution plot of ECOLI (Fig. 3a). This result is promising to further utilization of increasing cryo-EM entries in combination with the respective X-ray entries for obtaining more UMF data for the proteins that were not amenable to DSA.

In the normal scale main plot of YnaI (Fig. 6a), C_α_ pairs in the morphable parts are difficult to assess when we include all data points. On the other hand, C_α_ pairs in the unmorphable parts exhibit almost linear distance dependency. This type of main plot did not typically arise from a set of X-ray structures, but could be seen for a polypeptide consisting of a larger multi-chain complex, such as human hist1h2ab (histone H2A.2) as demonstrated previously^1^. The updated main plot, adding eight entries (chains) to the previous data, of hist1h2ab is shown in Fig. 6c.

**Fig. 6.**
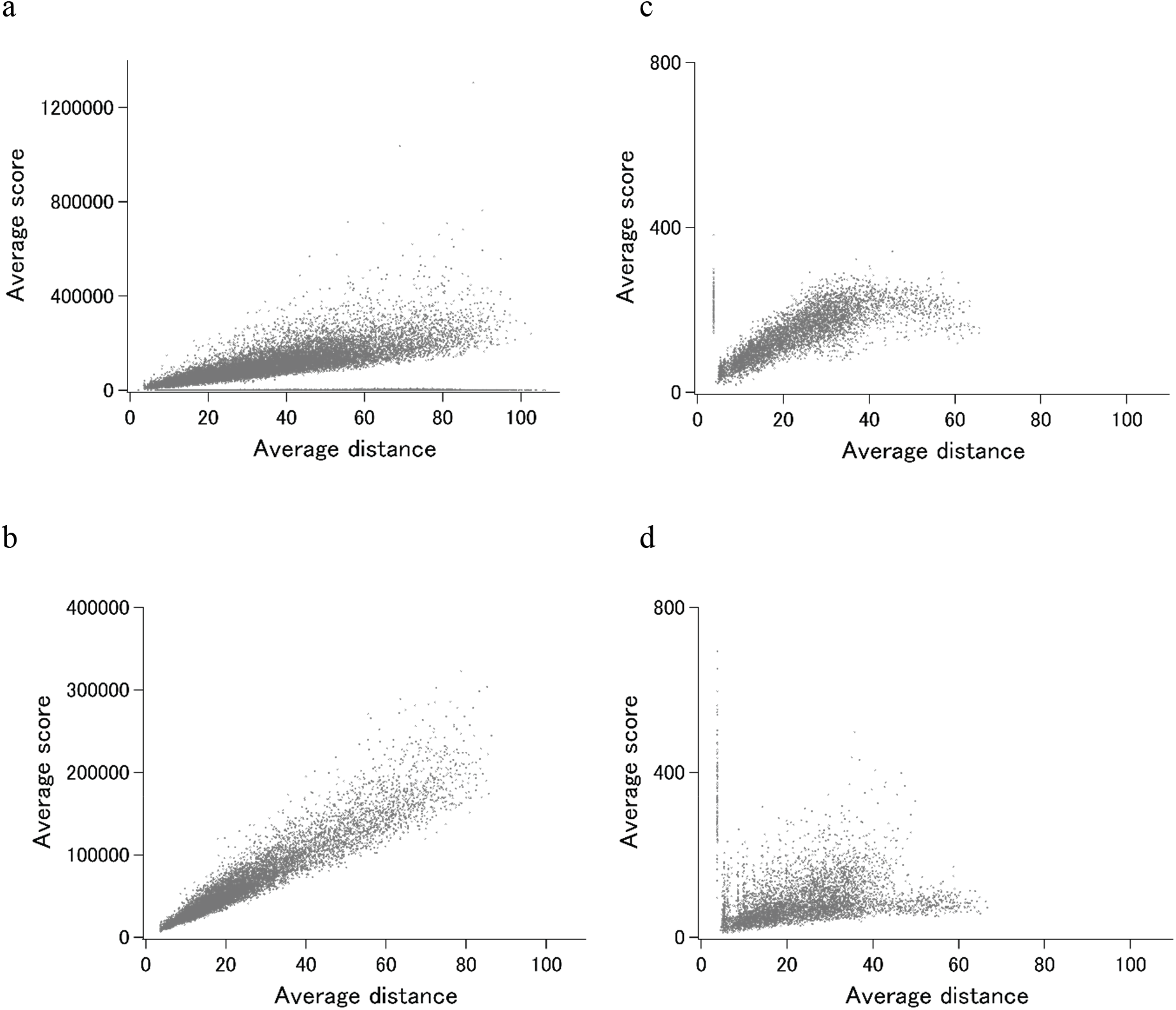
The main plot obtained from DSA. (a) YnaI of ECOLI (cryo-EM); (b) prepilin peptidase-dependent pilin of ECOLI (cryo-EM); (c) hist1h2ab of human (X-ray); and (d) hist1h2ab of human (cryo-EM).

For prepilin peptidase-dependent pilin (PPDD)^16^, which is another one of seven unusual eUMF value, the main plot (Fig. 6b) was found to be just like the pattern of hist1h2ab. These patterns are almost artificially reproducible by using values of model standard deviations for the average distances, and represent the examples of uniform C_α_ coordinate deviations at all positions in the polypeptide.

In Fig. 6d, the main plot of hist1h2ab by cryo-EM entries is shown. Although the used number of chains (22 chains, one chain per entries) are less than a half of X-ray entries (56 chains, one chain per entries), the cluster of C_α_ pairs by cryo-EM was found mostly in the lower average scores than that by X-ray data. Correspondingly, eUMF of 91.0 is lower than the xUMF of 143.5 which is substantially unchanged from the previous value with 48 chains usage. While the clear difference in the main plot pattern between cryo-EM and X-ray could arise from various factors in this case, our results suggest that cryo-EM is potentially effective in uncovering the full morphness range of a polypeptide with fewer structure determinations (PDB entries)

### Application to recent entries

How newly added PDB entries exhibit different backbone structures from that of previously deposited ones for the same protein is readily evaluated by the present method. Here we demonstrate an example obtained recently in July of 2019.

SOS response-associated protein YedK^17^ is included in the ECOLI reference proteome set and has a calculated xUMF of 321.7, using only three chains from two entries out of three entries deposited for UniProt ID of P76318. This value appeared to be possible to converge to a lower one, whereas even the progress plot could not be made with just one data point (number of chains = 3). The unused entry contains two regions of chain-break within the almost full-length model.

More recently, five entries were added to the PDB for the same protein but with different UniProt ID of A0A2S5ZH06 (100% identical to P76318). As the modeled length was the same as that of two entries for P76318, the new distance data can be added to the previous distance data, making the progress plot meaningful (Fig. 7a). The new data contributed to the data point 4 to 9, using 4 out of 5 entries and 6 chains from 4 entries (three chains from one entry 6KBZ). The average score value at the data point 9 translates to xUMF of 161.4, which is about a half of the previous 321.7. We have been making clear separation of entry usage and chain usage for most of over 5500 protein analysis, and the progress plot in this case is rearranged to 6 entry usage first followed by the redundant chain usage as shown in Fig. 7c where a redundantly used chain for P76318 is moved from data point 3 to 7. Although such a data switch in the progress plot did not cause any significant change in this case, data order switch would be useful for other proteins to see the impact (discontinuous drop of the average score) of some specific entries/chains in a set of many entries.

**Fig. 7.**
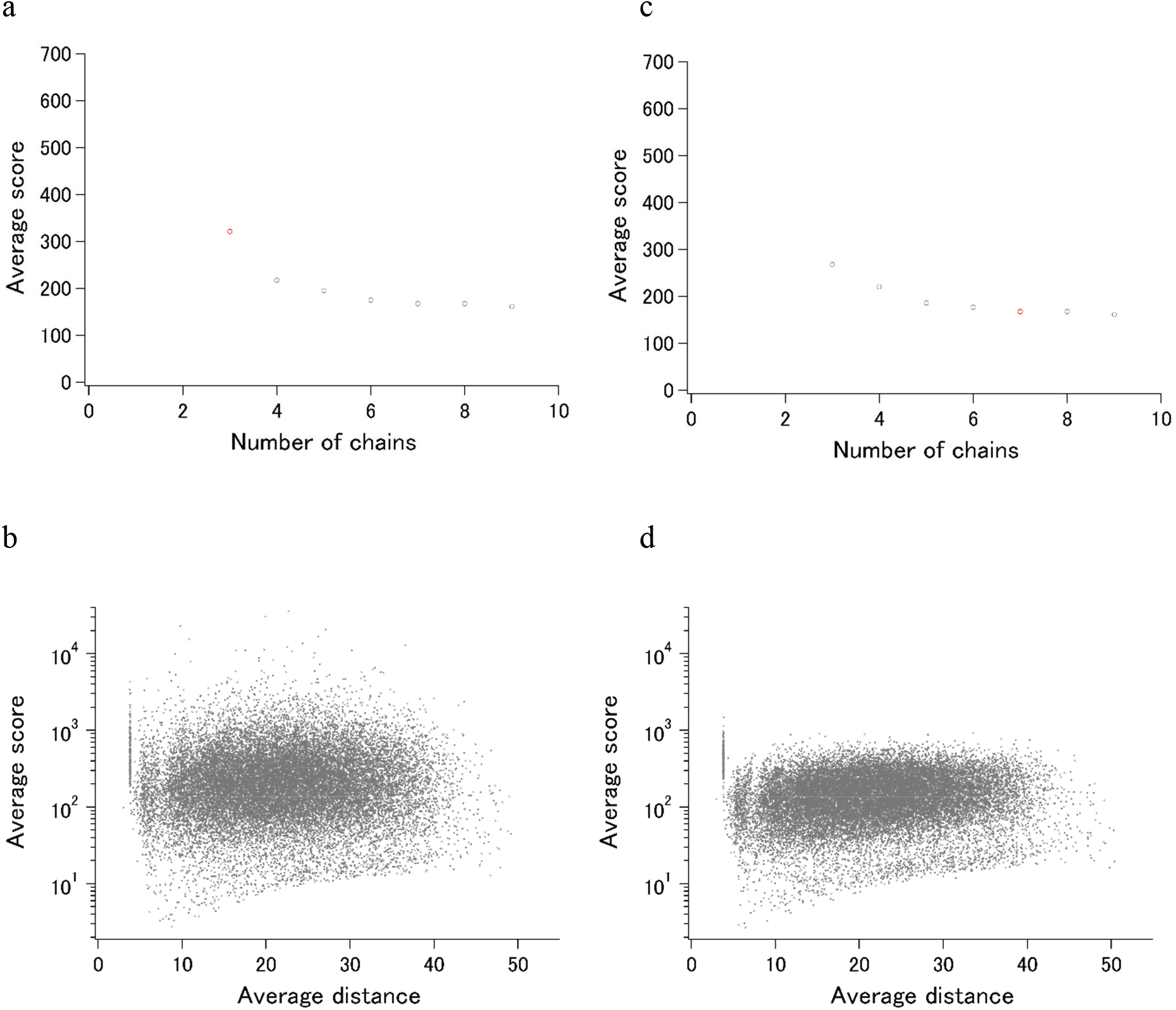
Recent examples of DSA update. (a) Progress plot of YedK (P76318/A0A2S5ZH06, the data point of only P76318 is shown in red.); (b) Progress plot of YedK (P76318/A0A2S5ZH06, the third data point in (a) is moved to the 7^th^ position); (c) semi-log main plot of YedK (P76318); (b) semi-log main plot of YedK (P76318/A0A2S5ZH06).

Two semi-log plots, one for P76318 and the other for P76318/A0A2S5ZH06, demonstrate the effect of entry/chain addition (Fig. 7b, d). Most of the C_α_-C_α_ pairs that appeared in the average score of over 1000, moved to much lower score region, whereas the highly variable pairs with low scores of less than 10 remain largely unchanged. This indicated that the first two entries (three chains) for YedK differed significantly from each other in the high morphness regions.

### Perspectives

The growth in the structural data accumulation has been outstanding for human proteins and the trend will undoubtedly continue in the future. Uncovering the morphness repertoire for any eukaryotic organism is very challenging due to the presence of splice variants and intrinsically disordered proteins. The total number of PDB entries for *E. coli* proteins is the second largest (after human proteins) and is exceptionally large among prokaryotic organisms. The present study shows that about a quarter of *E. coli*’s proteome set could have already been the subject of detailed morphness analysis. More coverage would be possible if we further include very similar orthologues from other bacteria such as salmonella species. UMF calculation by Cryo-EM data only (eUMF) was found to be unreliable in some cases but the data would undoubtedly be useful especially in combination with the corresponding X-ray data. It should be noted that we have not applied any resolution cut criteria for cryo-EM data analysis. While NMR data usage has not been given priority in our studies so far, it will have to be done in the future.

All DSA results can improve on a weekly basis in accordance with updates of the PDB contents. Importantly, the results presented here, when the sharing environment is established, can conform to FAIR data principles^18^ since no assumptions are involved in the analysis. Anyone can contribute to improve the quality of the data set by, for example, rationally changing the selection of the entries/chains based on their expertize of specific proteins.

## Methods

### Protein lists

ECOLI and HUMAN reference proteome set obtained from UniProt were initially used to search for the corresponding PDB entries to each of the UniProt IDs. For ECOLI, all UniProt IDs were checked to see how many PDB entries could be found for each protein, and then the protein list was sorted by amount of entries, showing that about 30% of the ECOLI proteins have at least one X-ray entries, and roughly a half of those have more than three entries. Drill down search by organism in the PDB was also used to supplement the protein list of *E. coli* outside the ECOLI proteome set. In fact, about 30% of *E. coli* PDB entries are of the UniProt IDs not found in the ECOLI set. These entries are used if they are of the structures of virtually identical sequences to any of the ECOLI set proteins. Thus, some of 980 xUMFs of ECOLI set are calculated from multiple UniProt IDs of almost identical sequences. For HUMAN, drill down search list was primarily used after UniProt ID sorting to find out which proteins are rich in the PDB entries. Roughly 23% (~4700) of HUMAN reference set and reviewed proteins were found to have at least one X-ray PDB entry. This means that the number of targets amenable for DSA is fairly more than the presently performed ~1000 if we carry out the analysis as extensively as has been done for ECOLI set.

### Archiving

Coordinate directories and data directories for proteins are separately made per UniProt ID under the corresponding organism top directory. Each coordinate directory contains the raw pdb files assigned to the respective UniProt ID. The data directory of each protein holds the following basic file set: (1) list file, containing information of PDB entries (chain IDs, space group, lattice constants, resolution, etc), the polypeptide range (initial and final sequence positions) selected for DSA, and the order of entry/chain usage in the progress plot (Fig. 7a,b). (2) Distance file, containing all C_α_ pair position numbers and the corresponding distances. (3) Score file, containing all C_α_ pair position numbers and the corresponding average distances, stdevs, scores, maximum and minimum distances. (4) Main plot scatter chart (and/or semi-log main plot scatter chart). (5) Progress plot table, and (6) progress plot scatter chart.

To date, over 200 species or group of species (~80 eukaryotic, ~110 bacterial, ~20 archaeal, and ~15 viral) have been archived and the total number of proteins analyzed so far exceeds 5500, although the degree of DSA completeness varies among the species. Completeness means how many multiple chains were included per entry. For example, our previous analysis of 300 human proteins used only one chain per entry because we selected the proteins having at least four entries. Although multiple chain usage from an entry do not always affect the UMFs substantially, there are some cases where many chain As from distinct entries are very similar to each other but chains A and B differs from each other. Thus, some of the xUMFs of human proteins would exhibit lower values when our chain usage completeness becomes higher.

The tables associated with the present study will be freely accessible from our web site (gses.jp).

### Data processing

DSA procedure was performed the same as previously described^1^. A Python script score-analyzer v22 has been modified from the previous version. Among many minor changes, such as the output function of a semi-log main plot chart, for example, up to 1,000,000 C_α_-C_α_ pairs (over 1400 amino acid length) are now can be processed, covering almost all of the currently deposited and continuously modeled structures. The longest polypeptide analyzed so far is the human U5 small nuclear ribonucleoprotein 200 kDa helicase (1397 residues, 65.4% of the full length is continuously modeled).

The 7TM structures for CNR1, a G protein-coupled receptor, were obtained from our archive at gses.jp/7tmsp. Each of the chains contains 200 amino acids, defined previously^19^ for reproducible comparison of all receptors including non-rhodopsin-like GPCRs.

### Mapping of UMFs to the proteome set

The protein list of ECOLI and HUMAN reference (reviewed) sets obtained from UniProt were used as a template. Sequence lists (FASTA) of these reference sets were used for the online analysis by SCAMPI to assign probable membrane proteins and the result was merged to the protein template list. The coverage of the present UMFs in the ECOLI set was evaluated first by trying to map all 1178 *E. coli* xUMFs to any of the proteins on the template list, resulting in 980 non-redundant assignments. For cryo-EM data, 56 out of 57 eUMF were assigned to the proteins on the template list, including 31 redundant assignment with the xUMF. In many cases of these 31 proteins, eUMF was found to be substantially lower than the corresponding xUMF, regardless of the number of chains used for analysis. When we mapped all 1039 xUMFs in the HUMAN reviewed reference set, no redundancy was found.

## References

1. Anzai, R. et al. Evaluation of variability in high-resolution protein structures by global distance scoring. Heliyon 4, e00510 (2018).

2. Burley, S. K. et al. Protein data bank (PDB): the single global macromolecular structure archive. Methods Mol. Biol. 1607, 627–641 (2017).

3. Okada, T. Protein structures and the nature of life. Science Trends (2018). at <https://sciencetrends.com>

4. The UniProt Consortium. UniProt: the universal protein knowledgebase. Nucleic Acids Res. 45, D158–D169 (2017).

5. Peters, C., Tsirigos, K. D., Shu, N. & Elofsson, A. Improved topology prediction using the terminal hydrophobic helices rule. Bioinformatics 32, 1158–1162 (2016).

6. Dunker, A. K. et al. Intrinsically disordered protein. J Mol Graph Model 19, 26–59 (2001).

7. Ahrens, J. B., Nunez-Castilla, J. & Siltberg-Liberles, J. Evolution of intrinsic disorder in eukaryotic proteins. Cell Mol. Life Sci. 74, 3163–3174 (2017).

8. Nogales, E. The development of cryo-EM into a mainstream” structural biology technique. Nat. Methods 13, 24–27 (2016).

9. Bai, X., McMullan, G. & Scheres, S. H. W. How cryo-EM is revolutionizing structural biology. Trends Biochem. Sci. 40, 49–57 (2015).

10. Raunser, S. Cryo-EM Revolutionizes the Structure Determination of Biomolecules. Angew. Chem. Int. Ed. Engl. 56, 16450–16452 (2017).

11. Yu, J. et al. A binding-block ion selective mechanism revealed by a Na/K selective channel. Protein Cell 9, 1–11 (2017).

12. Böttcher, B., Prazak, V., Rasmussen, A., Black, S. S. & Rasmussen, T. The Structure of YnaI Implies Structural and Mechanistic Conservation in the MscS Family of Mechanosensitive Channels. Structure 23, 1705–1714 (2015).

13. Steinbacher, S., Bass, R., Strop, P. & Rees, D. C. Structures of the prokaryotic mechanosensitive channels MscL and MscS. Curr Top Membr (2007).

14. Hua, T. et al. Crystal structures of agonist-bound human cannabinoid receptor CB1. Nature 547, 468–471 (2017).

15. Beckert, B. et al. Structure of a hibernating 100S ribosome reveals an inactive conformation of the ribosomal protein S1. Nat. Microbiol. 3, 1115–1121 (2018).

16. Bardiaux, B. et al. Structure and Assembly of the Enterohemorrhagic Escherichia coli Type 4 Pilus. Structure 27, 1082–1093.e5 (2019).

17. Thompson, P. S., Amidon, K. M., Mohni, K. N., Cortez, D. & Eichman, B. F. Protection of abasic sites during DNA replication by a stable thiazolidine protein-DNA cross-link. Nat. Struct. Mol. Biol. (2019). doi:10.1038/s41594-019-0255-5

18. Wilkinson, M. D. et al. The FAIR Guiding Principles for scientific data management and stewardship. Sci. Data 3, 160018 (2016).

19. Okada, T. Comparative analysis of the heptahelical transmembrane bundles of G protein-coupled receptors. PLoS One 7, e35802 (2012).

